# Crosstalk between Rac and Rap GTPases in migrating cells

**DOI:** 10.1101/2025.02.14.637886

**Authors:** Ricarda Lüttig, Suchet Nanda, Haritha T. Chandran, Leif Dehmelt

## Abstract

To enable effective cell migration, local cell protrusion has to be coordinated with local cell attachment. Here, we investigate spatio-temporal activity patterns of key regulators of cell protrusion and adhesion, the small GTPases Rac and Rap, in migrating cells. These analyses show that Rac activity correlates very tightly with instantaneous cell protrusion events, while the Rap activity stays elevated for prolonged time periods after protrusion and is also detectable before cell protrusion. Direct analysis of activity crosstalk in living cells via light-based perturbation methods revealed that Rap can efficiently activate Rac, however, reciprocal crosstalk from Rac to Rap was not detectable. These findings suggest that Rap plays an instructive role in the generation of cell protrusions by its ability to activate Rac. Furthermore, prolonged Rap activity suggests that this molecule also plays a role in maintenance or stabilization of cell protrusions. Indeed, morphological analysis of Rap1-depleted A431 cells revealed a significant reduction of the cell attachment area, suggesting that Rap stimulated cell adhesion might indeed stabilize newly formed protrusions. Taken together, our study suggests a mechanism, by which cell protrusion is coupled to cell adhesion via unidirectional crosstalk that connects the activity of the small GTPases Rap and Rac.

## Introduction

Directed cell migration plays a fundamental role in many biological processes including embryonic development, the immune response and regeneration of injured tissues (Trepat et al., 2012). In addition to these physiological roles, cell migration can also drive the progression of diseases, such as the aberrant migratory behavior of malignant cancer cells that drives metastasis and cancer invasion (Scarpa & Mayor, 2016; Trepat et al., 2012).

The process of cell migration involves four steps. 1) First, cell protrusions are formed at the front of migrating cells. 2) Subsequently, new adhesions are generated at the leading edge of the cell. 3) Contractile structures, called stress fibers are formed that are linked to these adhesions, and that generate forces between the front and back of the cell. 4) Finally, the trailing end of the cell detaches and retracts after adhesions at the back of the cell are released. This last step generates excess dorsal surface to enable newly generated protrusions at the cell front (Ridley, 2001).

To facilitate migration, the processes of cell protrusion, adhesion, and retraction must be precisely coordinated in space and time. Consequently, the signaling network components that regulate these processes have to be closely interconnected. Interestingly, recent studies have shown that the network components that control various aspects of cell migration are controlled by positive and negative feedback loops that generate excitable system dynamics (Bement et al., 2015; Devreotes et al., 2017; Graessl et al., 2017; Miao et al., 2017; Yang et al., 2016). The causal links that connect protrusion and retraction signals can either enable highly dynamic cell shape changes, resulting in a more random, exploratory migration mode with frequent changes in direction (Arrieumerlou & Meyer, 2005; Nanda et al., 2023), or result in a more persistent directional migration mode via a more stable spatial segregation of protrusion and retraction (Patwardhan et al., 2024; Svitkina et al., 1997). For the spatio-temporal coordination of the many processes that occur during cell migration, regulators of the Rho and Ras families were shown to play important roles (Devreotes et al., 2017).

Rac1, a member of the Rho family of GTPases, is a master regulator for the formation of flat cell protrusions, which are called lamellipodia (Ridley et al., 1992). Within these structures, Rac1 activates the WAVE complex, a regulatory molecule that subsequently stimulates the actin nucleator Arp2/3. The newly formed actin filaments generate a polymerization-driven force that pushes the cell edge forward (Ridley, 2001; Steffen et al., 2013). After new adhesions are formed, another Rho family GTPase called RhoA stimulates contractile forces, which are generated by the activity of the molecular motor Myosin-II on actin filaments (Ridley, 2001).

The Ras/Rap family of small GTPases is also thought to play important roles in cell migration (Devreotes et al., 2017; Wittchen et al., 2005). Similar to other small GTPases, Ras/Rap proteins are activated by particular guanine nucleotide exchange factors (GEFs), inhibited by specific GTPase activating proteins (GAPs) and they relay their activity state to cellular processes via specific effectors. Interestingly, previous studies suggested that the activity of Ras/Rap proteins might be linked to other regulators of cell migration, in particular, to the cell protrusion master regulator Rac1. Based on previous studies, Ras/Rap proteins can act upstream of Rac1, for example via the GEF Tiam1 (Lambert et al., 2002) and also downstream of Rac1 via ROS (Diekmann et al., 1994; Ferro et al., 2014) or growth factor signaling (Joshi et al., 2023). Ras GTPases are well-studied oncogenes that mediate growth factor signaling (Hobbs et al., 2016). Rap GTPases are key regulators of integrin-mediated cell matrix adhesion and cadherin-mediated cell-cell-junction formation. In particular, the Rap1 isoforms are known to activate integrins via the Rap1-GTP-interacting adapter molecule (RIAM) (Lafuente et al., 2004).

Here, we developed and applied new, improved live cell activity sensors to investigate the function of Ras/Rap proteins in the keratinocyte-derived cancer cell line A431. Our studies show that the activity of Rap-type GTPases is strongly enriched at the leading edge of migrating cells. In contrast, we find that the activity of the closely related Ras-type GTPases show very little local enrichment during directional migration. Interestingly, while Rac activity is exclusively enriched only during time frames of active cell protrusion, Rap activity is already observed before significant activation of Rac and then maintained for a longer time after the cell protrusion event. These complex dynamics raised the question, if Rap acts upstream or downstream of Rac in these cells. Direct, light-controlled activity manipulation revealed that the specific Rap isoform Rap1a acts upstream of Rac1, but conversely, that Rac1 does not activate Rap-related GTPases. Taken together, our findings reveal a clear hierarchy between Rap and Rac activities and support a model, in which Rap signals are necessary and permissive for Rac-dependent cell protrusion events, and for the subsequent processes that consolidate newly formed cell protrusions.

## Results

### Ras/Rap activity in migrating cells

In previous studies of cell migration in the keratinocyte derived A431 cell line, we found that highly dynamic protrusion-retraction cycles are stimulated by an unexpected crosstalk that activates the cell contraction regulator Rho downstream of the cell protrusion regulator Rac1 (Nanda et al., 2023). These studies raised the question, which signals act upstream of Rac to initiate the dynamic protrusion-retraction cycle. Previous studies, in particular in *Dictyostelium*, suggested that members of the Ras/Rap family of small GTPases act upstream of Rac1, however, little information was available on the spatio-temporal activity state of these molecules in mammalian cells. To fill this gap in our knowledge, we used a similar strategy as in our previous study (Nanda et al., 2023) to generate sensitive activity sensors for Ras and Rap-family GTPases (Figure 1a). Briefly, we fused two tandem copies of effector domains that either preferentially bind the active form of Rap-subfamily (RalGDS-GBD) or Ras subfamily (Raf-GBD) GTPases to a fluorescent protein, expressed the sensor proteins at very low levels using the delCMV promotor (Watanabe & Mitchison, 2002) to avoid competition with effector proteins, and monitored the plasma membrane translocation of these sensors via total internal reflection fluorescence microscopy (TIRF-M). Using these sensors, we observed distinct dynamic patterns of Rap activity in A431 cells, including transient pulses in central cell attachment areas and both transient and more prolonged activity at the cell edge that was closely correlated with active cell protrusion (Figure 1b-e; Movies 1 and 2). Active Ras on the other hand was more homogenously distributed and did not show strong local enrichment (Figure 1b-d; Movie 1), which is in agreement with its major role in controlling cell proliferation and cell growth.

**Fig. 1.**
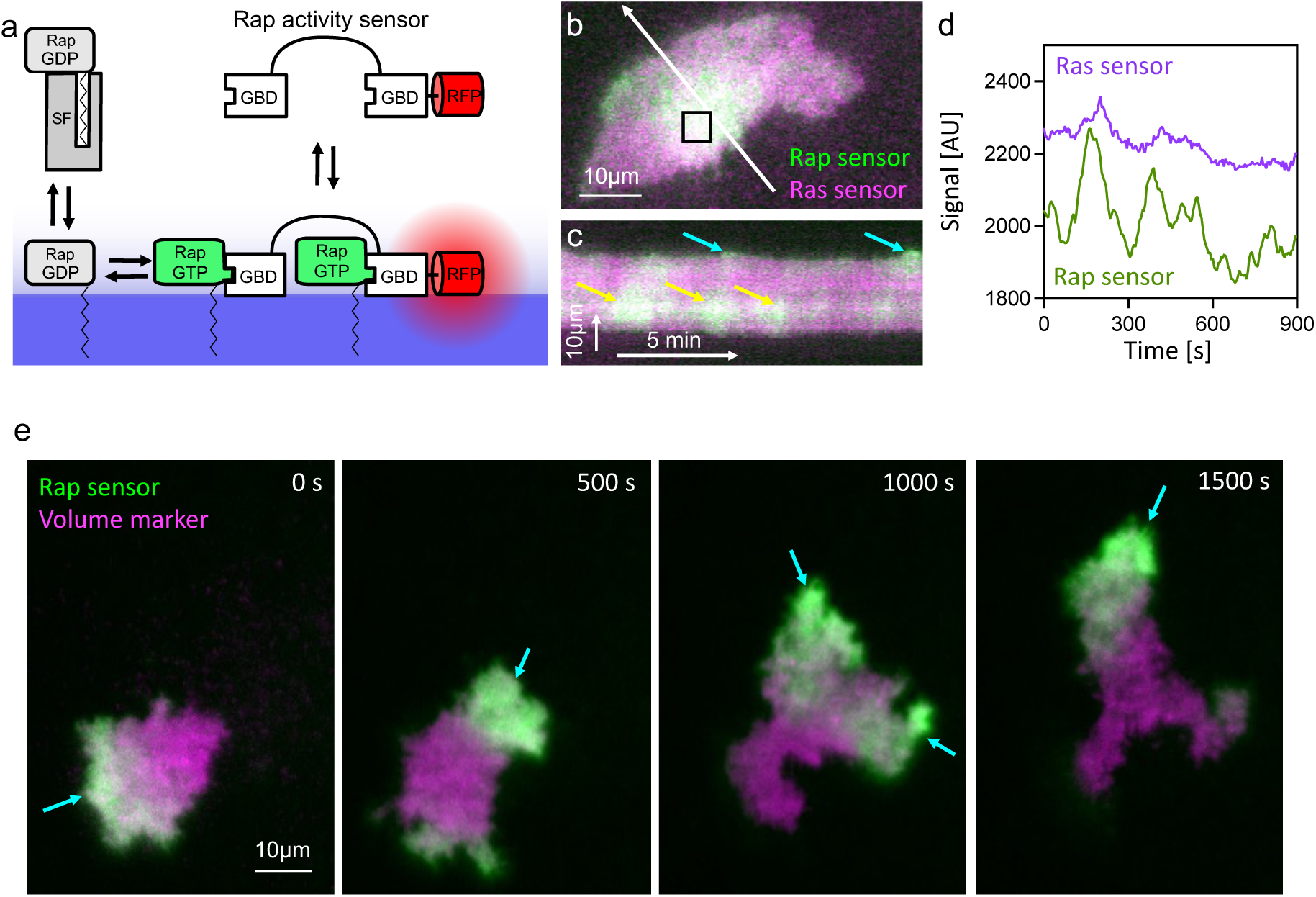
Spatio-temporal Ras and Rap activity dynamics. **a** Schematic for Ras and Rap activity sensors used in this study. Similar to other GTPases, inactive GDP-bound Ras/Rap is largely cytosolic, where it binds to a solubilization factor (SF). Guanine nucleotide exchange factors (GEFs) catalyze the transition into the active, GTP-bound Ras/Rap state, which is preferentially localized at the plasma membrane, where it interacts with effectors via GTPase binding domains (GBDs). The sensors employed in this study contain tandem repeats of GBDs that are fused to a fluorescent protein, and their translocation from the cytosol to the active GTPase at the plasma membrane is monitored via total internal reflection fluorescence microscopy (TIRF-M). **b** Representative TIRF image of a A431 cell co-expressing the Rap (mCitrine-2xRalGDS-GBD) and Ras (2xRaf-GBD-mCherry) activity sensors (see also Movie 1). Cyan arrows point at cell protrusions and yellow arrows point at Rap activity pulses in central cell attachment areas. **c** Kymograph corresponding to white arrow in **b**. **d** Quantification of signal intensity changes in the black square region in **b**. **e** Sequential TIRF images of a single migrating A431 cell that expresses the Rap activity sensor and a volume marker (see also Movie 2). Cyan arrows point to newly formed cell protrusions.

To quantify Rap and Ras activity dynamics, we analyzed how their signals change with protrusion and retraction of the cell edge using a modification of the ADAPT ImageJ plugin (Figure 2) (Barry et al., 2015; Nanda et al., 2023). These analyses show that Rap activity signals strongly and positively correlate with cell edge velocity (Figure 2d, e, m; Movie 3). On average, Rap activity is slightly delayed with a maximal correlation at ≈1min (65s) after cell protrusion.

**Fig. 2.**
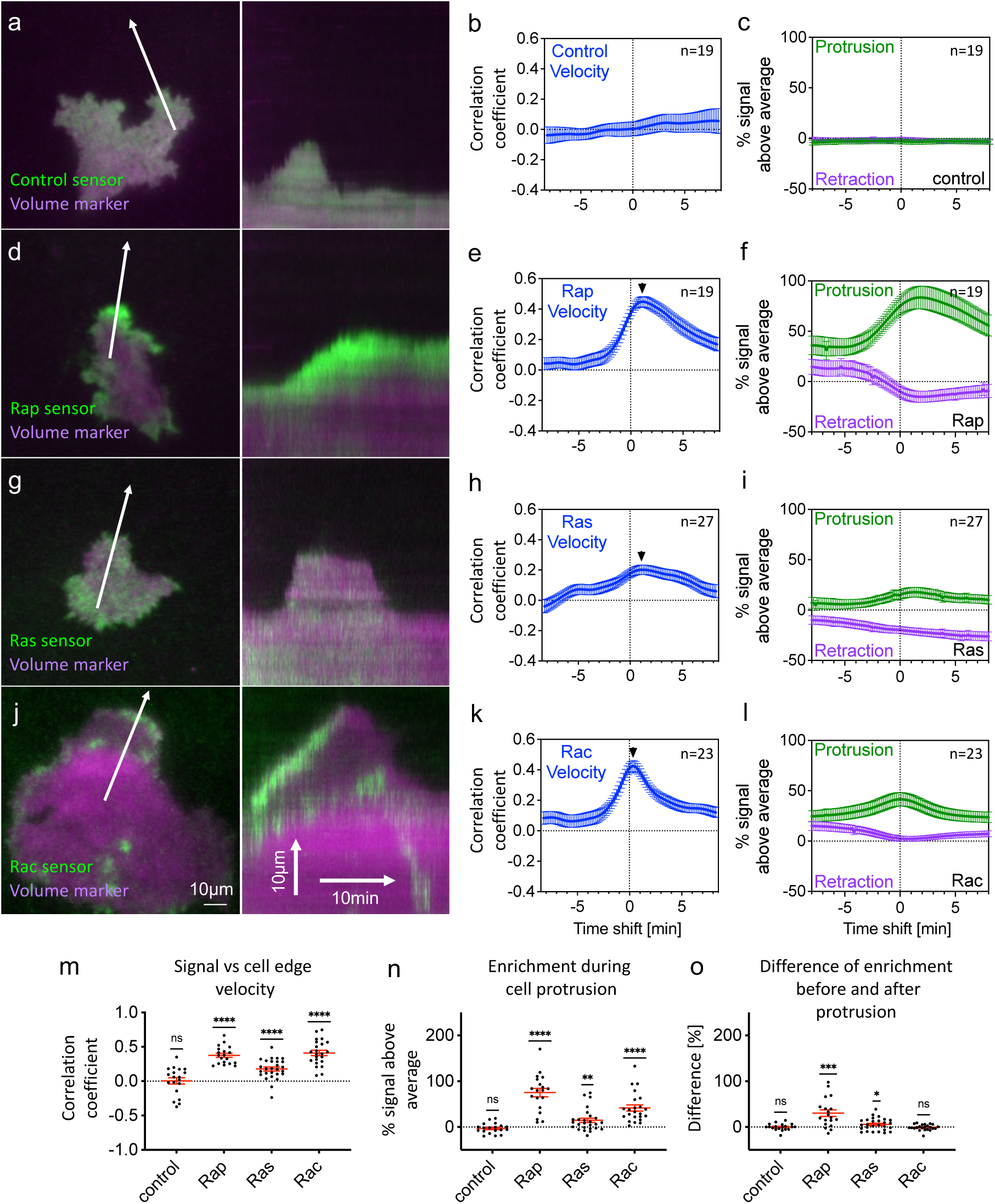
Ras/Rap activity dynamics in migrating cells. a-l. Analysis of control (a-c), Rap (d-f), Ras (g-i) and Rac (j-l) activity dynamics at the cell periphery during protrusion-retraction cycles (see also Movie 3) **a,d,g,j** Representative TIRF images of the activity sensor and a volume marker (left), and kymographs (right) corresponding to the white arrows in left panels**. b,e,h,k** Crosscorrelation functions for sensor signals and cell edge velocity plotted against the time shift between these measurements. Black arrow heads point to time shift with maximal signal after cell protrusion. **c,f,i,l** Enrichment of sensor signals in protrusions (> 0.5 µm/min) and retractions (< -0.5 µm/min). Values are normalized to mean control sensor measurements. **m** Measurements of the signal-cell edge velocity correlation coefficient at a time shift of 0 min of the correlation functions shown in **b,e,h,k**. **n** Measurements of the signal enrichment at a time shift of 0 min of the enrichment plots shown in **c,f,i,l**. **o** Difference between the signal enrichment before and after protrusion (defined as time shift of 0 min), extracted from data shown in **c,f,i,l**. Data corresponding to Rac activity shown in **j-l** and in the last column of **m-o** were reproduced from (Nanda et al., 2023a) (data is protected by CC BY4.0 licence). (*: P<0.05; **: P<0.01; ***: P<0.001; ****: P<0.0001; ns: not significant; One-sample, two-sided t-test; n>18 cells from at least 3 independent experiments, exact numbers of cells are indicated in panels **c,f,i,l**). Error bars represent the standard error of the mean (SEM).

The cell edge velocity correlation cannot distinguish between activity signal increase with positive velocity during cell protrusion or signal decrease with negative velocity during cell retraction. Furthermore, the cell edge velocity correlation only quantifies how similar signal changes are compared to cell edge velocity changes and does not indicate how strongly the sensor signals are locally enriched within cells. We therefore also performed a protrusion-retraction enrichment analysis that we recently developed (Nanda et al., 2023), to quantify how much the sensor signals are increased at the cell edge compared to the entire cell attachment area during cell protrusion and cell retraction phases. These analyses showed that Rap activity is strongly enriched by over 80% with a maximum at ≈2min (112s) after cell protrusion (Figure 2f, n). Conversely, Rap activity is depleted by about 15% at ≈2min (112s) after cell retraction. Correlation and enrichment measurements of signals of the Ras activity sensor were much weaker compared to the Rap activity sensor (Figure 2g-i, m-n; Movie 3), but were nevertheless clearly above levels obtained using a cell volume marker that served as control (Figure 2a-c, m-n; Movie 3).

Interestingly, the Rap correlation and enrichment measurements showed a strong asymmetry for time periods before and after cell protrusion. This was particularly clear for the signal enrichment during protrusion. We did not observe such an asymmetry using an identical assay for Rac activity in our previous studies (Nanda et al., 2023). For comparison, the measurements that were obtained in this previous study are reproduced in Figure 2j-l (see also Movie 3). To quantify this asymmetry, we subtracted the mean enrichment in the time period before protrusion from the mean enrichment after protrusion (Figure 2o). This quantification clearly shows that Rap activity is more strongly enriched at the leading edge of the cell after protrusion compared to before protrusion. No asymmetry was measurable for Rac activity, and Ras activity showed a weak, intermediate trend (Figure 2o).

### Crosstalk between Rap and Rac GTPases

The asymmetric enrichment of Rap activity relative to cell protrusion events suggests a delayed activation of this regulator. In contrast, the more symmetric enrichment of Rac activity suggests a tight spatio-temporal link to cell protrusion. Indeed, the maximal enrichment of Rap activity is substantially shifted after cell protrusion (τ=112s), while the maximal enrichment of Rac is not shifted at all (τ=0s). Intuitively, this delay could indicate that Rap acts downstream of Rac. Simultaneous imaging of both Rac and Rap activity supports this idea: While both Rap and Rac are highly active during cell protrusion, Rac activity quickly diminishes after protrusion, while Rap activity often remains high (Figure 3a; Movie 4). However, these experiments also revealed a more complex pattern. In addition, Rap activity was often observed before cell protrusion in the absence of active Rac (Figure 3a; Movie 4). Thus, we both observed events, in which Rap is active before Rac, and events in which Rac is active before Rap. As previous studies could explain both a causal link from Rac to Rap (Lambert et al., 2002), as well as from Rap to Rac (Diekmann et al., 1994; Ferro et al., 2014; Joshi et al., 2023), we directly investigated potentially bidirectional crosstalk between these molecules by combining rapid activity perturbations with readouts of the signal network response.

**Fig. 3.**
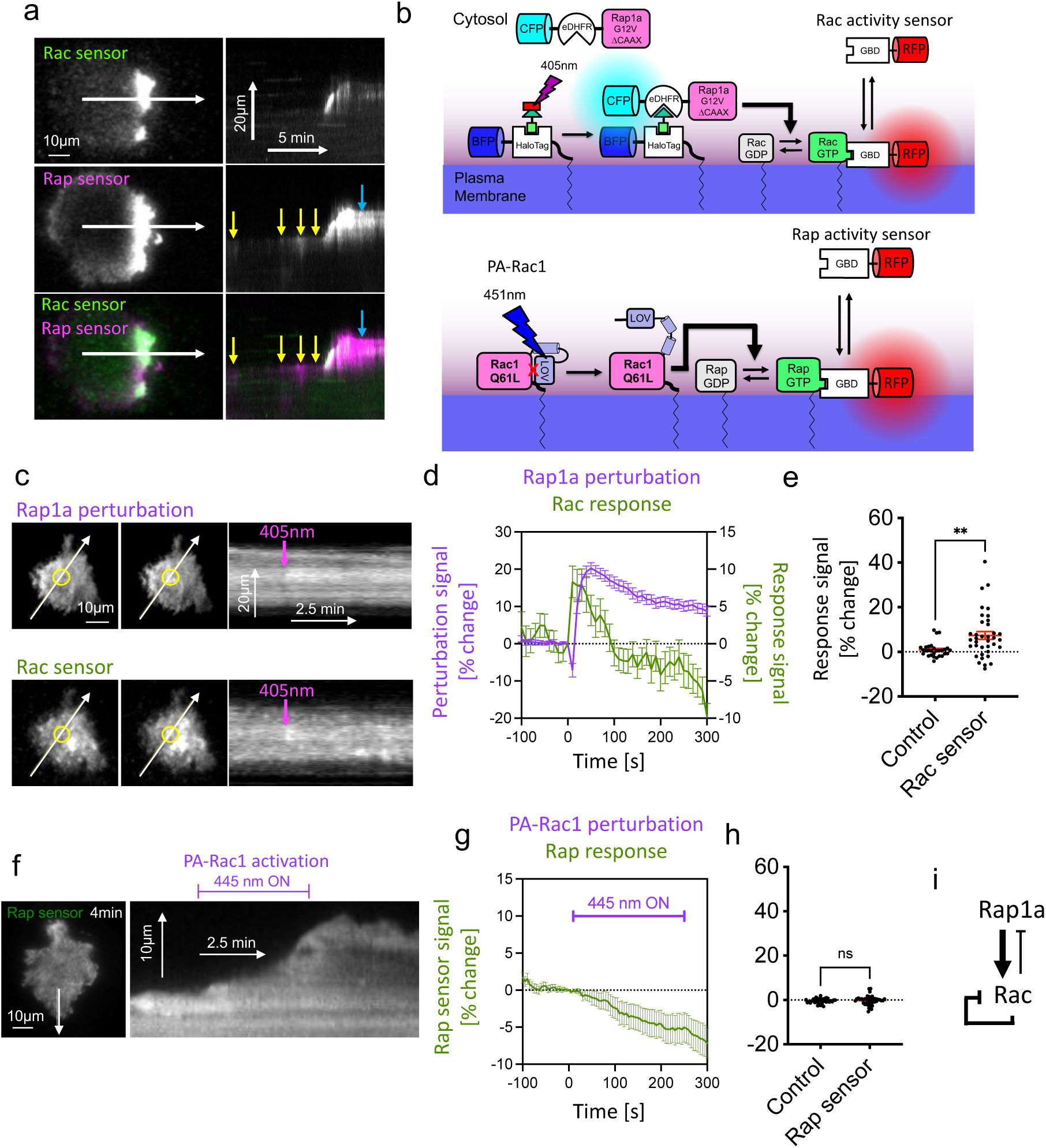
Investigation of GTPase crosstalk revealed unidirectional activation of Rac downstream of Rap. **a** Representative TIRF images (left) and corresponding kymographs (right) of A431 cells co-expressing Rap (mCitrine-2xRalGDS-GBD) and Rac (mCherry-3xp67phox-GBD) activity sensors (see also Movie 4). Yellow arrows point to weak transient Rap signals that can be detected before Rac activation. Blue arrows point to prolonged Rap activity after initial protrusion. **b** Schematic for the photochemical dimerizer-based perturbation method (top) and for PA-Rac1 mediated perturbations (bottom), and their combination with activity measurements. **c** Left: Representative TIRF images of an A431 cell that co-expresses the dimerization fusion proteins mTurquoise2-NES-eDHFR-Rap1a G12V (“Rap1a perturbation”), mTagBFP-Halotag-CAAX (not shown) as well as the Rac activity sensor mCherry-3xp67^phox^GBD (“Rac sensor”). Right: Kymographs that correspond to the white arrows in left panels (see also Movie 5). **d** Quantification of local Rap1A perturbation and parallel measurement of Rac activity sensor recruitment dynamics. **e** Quantification of the early Rac activity response (n=29 for control and n=37 cells for the experimental condition, from 3 independent experiments). The difference between the mean measurements of 5 timepoints before and after the onset of illumination is shown. **f** Left: Representative TIRF image of an A431 cell that co-expresses mCerulean-PA-Rac1 and the mCitrine-2xRalGDS-GBD Rap activity sensor. Right: Kymograph that corresponds to the white arrow in the left panel (see also Movie 6). **g** Quantification of Rap1A recruitment dynamics to the entire cell attachment area. **h** Quantification of the early Rap activity response. The difference between the mean measurements of 5 timepoints before and after the onset of illumination is shown (n=38 for control and n=42 cells for the experimental condition, from 3 independent experiments). **i** Proposed signal network that links Rap and Rac activity in cells. (**: P<0.01; ns: not significant; two-sided Student’s t-test). Error bars represent the standard error of the mean (SEM).

To test, if Rap can activate Rac, we used a method that we previously developed to introduce rapid perturbations in signal networks at the plasma membrane by photochemically-induced dimerization (Chen et al., 2017), (Figure 3b). Of the five known human Rap isoforms we concentrated our investigations on the well-studied Rap1 subfamily and specifically on Rap1a, as this isoform is expressed at a higher level in A431 cells compared to Rap1b (Klijn et al., 2014). Briefly, we fused a dominant-positive mutant of Rap1a (G12V) to a first heterodimerization domain: *E. coli* dihydrofolate reductase (eDHFR). In addition, we expressed a second heterodimerization domain, Halo-Tag, fused to the CAAX-Box plasma membrane targeting sequence. We treated cells with the photocaged dimerizer NvocTMP-Cl and induced plasma membrane targeting of Rap1a (G12V) by local photouncaging of the dimerizer. We combined this method with the Rac activity sensor that we also used in Figure 2j-l. Using this method, we measured a very rapid and transient Rac activity response at the uncaging spot (Figure 3c-e; Movie 5). These observations clearly show that Rap1a acts upstream of Rac in the A431 cell line. Interestingly, while the levels of active Rap1a remained elevated for more than 5 minutes, the activity of Rac decreased much faster and reached baseline levels already after 1.5 min. This suggests that the signal system that connects Rap and Rac activities also includes an effective and rapid negative regulation of Rac.

To investigate the reciprocal causality, if Rac can activate Rap, we used the previously established, PA-Rac1 construct (Wu et al., 2009), which enables light-controlled activation of Rac1 in living cells (Figure 3b), and we combined perturbations induced by this construct with Rap activity measurements via the activity sensor used in Figure 2e-f. As expected, rapid activation of Rac1 with light lead to the generation of new lamellipodial cell protrusions. However, in contrast to lamellipodia that spontaneously form in A431 cells, which always enriched high levels of active Rap (Figure 1e, and Figure 2d), the PA-Rac1 triggered lamellipodia did not display an enrichment in Rap activity (Figure 3f; Movie 6). Furthermore, quantification of the Rap sensor signal in the entire cell attachment area did not show any Rap activation by Rac, and instead even suggested a small reduction of its activity (Figure 3g-h).

Taken together these two observations suggest a largely unidirectional causality between Rap and Rac activities (Figure 3i): Rap can very quickly and efficiently activate Rac within seconds. Subsequently, Rac activity is inhibited, presumably via negative feedback regulation that acts within 1-2 minutes. Conversely, Rac does not strongly influence Rap activity. The weak reduction of Rap activity downstream of Rac over the time course of about 5 minutes is too slow to explain negative feedback regulation of Rac1 by itself, but it might play a role in slower processes that act in parallel.

### Efficient cell attachment is dependent on the Rap1a isoform

Parallel imaging of Rac and Rap activity showed that these regulators are both activated very strongly during active protrusion (Figure 3a). However, as shown in Figure 3a and in Figure 2d-f, o, Rap activity remained elevated much longer, even after the instantaneous protrusion events. As new cell attachment sites have to be established during and shortly after cell protrusion, this observation fits to the well-known role of Rap1 in stimulating new cell adhesions (Boettner & Van Aelst, 2009). To investigate, if these processes are indeed controlled by Rap1 GTPases in A431 cells, we combined knock-down experiments with phenotypic readouts. Our studies reveal that the knockdown of Rap1a leads to a significant reduction of cell adhesion area, suggesting that this molecule is required for normal adhesion and spreading of A431 cells on extracellular substrates (Figure 4a-b).

**Fig. 4.**
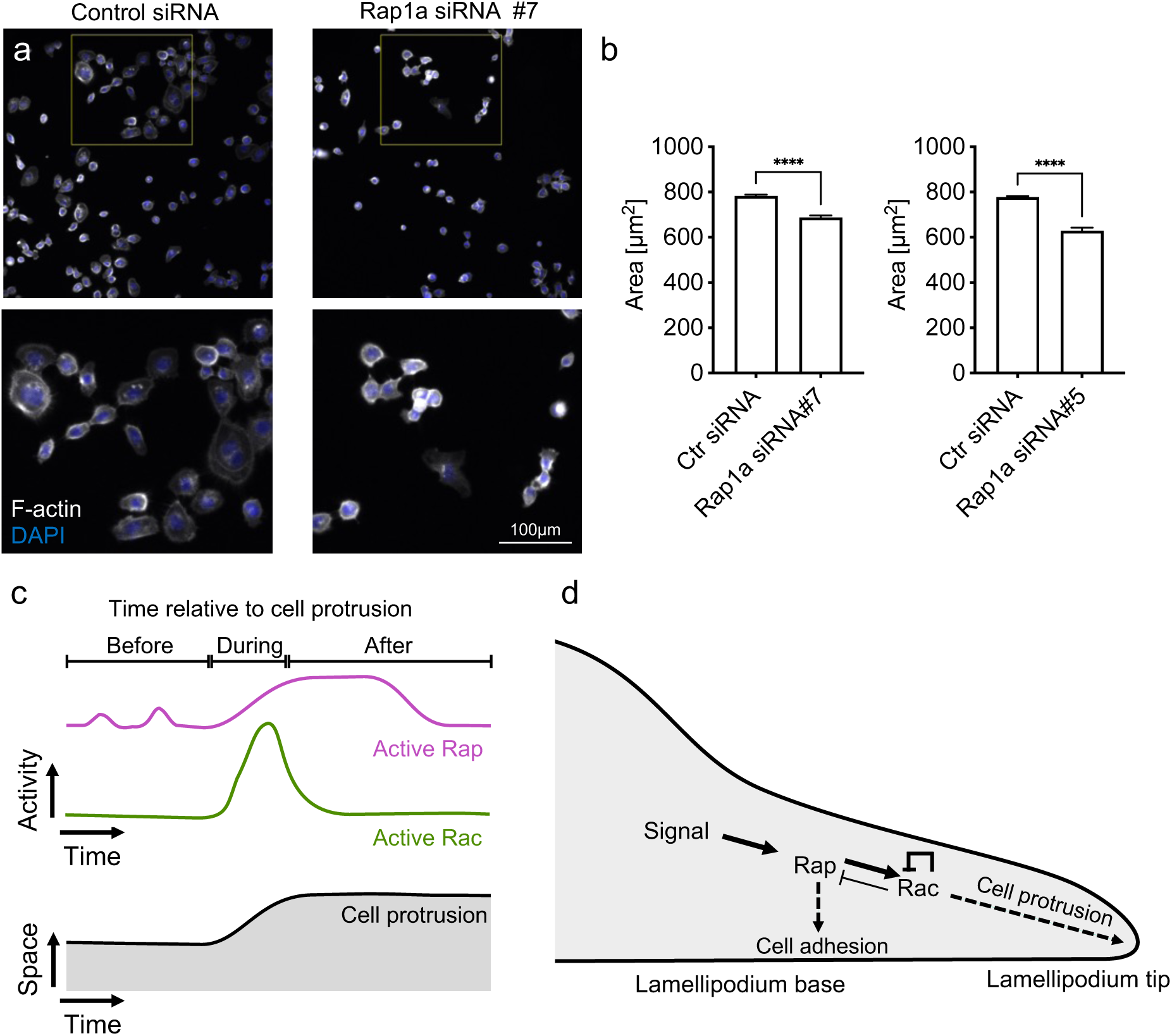
Efficient cell attachment is dependent on the Rap1a isoform. **a**-**b** Effect of Rap1a knockdown on the cell attachment area. Cells were stained for DNA and filamentous actin; area quantification was performed using Cell Profiler. Knockdown efficiency was highly efficient (91±3% for siRNA#7;n=3 and 93.4±1% for siRNA#5; n=4; mean±SEM). **c** Schematic for the activity state of Rap and Rac in relation to cell protrusion. **d** Schematic for the proposed role of Rap in cell protrusion. An upstream signal, for example from growth factor receptor activity, stimulates Rap, which in turn activates Rac, leading to cell protrusion. Rac can quickly inhibit itself, for example by activating Rho (Nanda et al., 2023), and thereby limit cell protrusion. Rac might be able to also inhibit Rap, however, this potentially inhibitory role is relatively weak and/or slow. Rap activity can therefore persist over longer periods of time to stabilize the newly formed protrusion by the formation of new cell adhesions. (****: P<0.0001; two-sided Student’s t-test; left: n=13578 for control and n=8539 cells for siRNA #7, from 3 independent experiments; right: n=20375 for control and n=11920 cells for siRNA #5, from 4 independent experiments). Error bars represent the SEM.

Taken together our studies show that Rap and Rac activities are coordinated in space and time during migration of A431 cells (Figure 4c-d). We reveal a clear hierarchy between these signal molecules and find that the isoform Rap1a can efficiently activate Rac. We also find that Rac is downregulated in the continued presence of active Rap. This suggests negative feedback regulation of Rac that is not mediated by Rap, but instead by another, Rap-independent mechanism (Figure 3i, 4d).

These direct investigations of activity crosstalk can fully explain the observed coordination between Rap and Rac activities during cell protrusion-retraction cycles (Figure 4c): Initial Rap activity is already observed before Rac activity, presumably due to upstream signals, for example downstream of growth factor receptor signaling. If Rap is sufficiently active, it activates Rac, leading to efficient cell protrusion (Figure 4d). Rac activity is inhibited via negative feedback, for example via activation of Rho (Nanda et al., 2023) and subsequent activation of Rac GAPs (Guilluy et al., 2011), while Rap activity is still high, supporting subsequent steps, including cell adhesion (Figure 4d). Taken together, our investigations suggest a mechanism that ensured effective cell protrusion by coupling this process to increased cell adhesion in space and time via unidirectional crosstalk from Rap1 to Rac1.

## Materials and Methods

### Cell culture

A431 (CRL-1555, ATCC) cells were maintained using standard cell culture techniques at 37°C and 5% CO2 using DMEM medium (2 mM L-Glutamine, PAN Biotech and 10% FBS, PAN Biotech or Sigma-Aldrich Chemie GmbH/Merck). For imaging, cells were either plated onto glass-bottom dishes (MatTek) or LabTek glass surface slides (Thermo Fischer Scientific), which were coated with 10 μg/ml Fibronectin for 45 min at RT. Transfection of plasmid DNA was performed using Lipofectamine^TM^ 3000 (Invitrogen, Thermo Fisher Scientific).

### siRNA mediated knockdown

Knockdown experiments were performed using ON-TARGETplus siRNAs (Dharmacon^TM^, Horizon Discovery): (siControl: #2 5′-UGGUUUACAUGUUGUGUGA-3’, Rap 1A: #5 5’-GAACAGAUUUUACGGGUUA-3’, #7 5’-GCGAGUAGUUGG CAAAGAG-3’). For knockdown, A431 cells were transfected with 30 nM siRNA using Lipofectamine^TM^ RNAiMAX (Invitrogen, Thermo Fisher Scientific) and incubated for 24h, followed by cell passaging to remove excess siRNAs. Knockdown efficiency was assessed 48h after siRNA treatment via qPCR. Briefly, RNA samples were prepared using the RNeasy^®^ Mini Kit (Qiagen) and QIAshredder (Qiagen) and processed by subsequent cDNA synthesis (GoScript^TM^ Reverse Transcription System, Promega), followed by qPCR. 96h after siRNA transfection, phenotypic analysis was performed on fixed cells.

### Plasmid constructs

The Rac activity sensor construct, delCMV-mCherry-3x-p67phox-GBD, and the control constructs delCMV-mCitrine and delCMV-mCherry were described previously (Nanda et al., 2023). The PA-Rac construct used for optogenetic perturbations, mCerulean-PA-Rac1Q61L, was a kind gift from Klaus Hahn (University of North Carolina) (Wu et al., 2009). The plasma membrane associated HaloTag construct used for photochemically-induced dimerization, mTagBFP-HaloTag-CAAX, and the photo-caged, small molecule dimerizer, NvocTMP-Cl, were described previously (Chen et al., 2017). The Rap activity sensor construct delCMV-mCitrine-2xRalGDS-GBD was generated by restriction digestion and ligation of 3xGFP-RalGDSRBD (kind gift from Philippe Bastiaens, MPI Dortmund) and delCMV-mCitrine-Rhotekin-GBD (Kamps et al., 2020), using AccIII and MfeI, and subsequent doubling of the RalGDS-GBD insert by Gibson assembly using AccIII and the primers 5’- CATGGACGAGCTGTACAAGTCCGGAGGTTCCGGAAGTGGATCCGCGCTGCCGCTCTACAAC-3’ and 5’-GCGCAGCTCGAGATCTGAGTCCGGATCCACTTCCGGAACCGGTCCGCTTCTTGAGGAC-3’. The Ras sensor delCMV-2xRaf-GBD-mCherry was cloned by restriction digestion and ligation of RafRBD-mCherry (kind gift from Philippe Bastiaens, MPI Dortmund) and delCMV-mCherry using NheI and AgeI, followed by doubling of the Raf-GBD via Gibson assembly using AgeI and the primers 5’- ACGGGTTCTGGAAGTGGATCGGTTCTCATGTCCCTGGTGGAGGC-3’ and 5’- GGCCTCCACCAGGGACATGGAATTCGATCCACTTCCAGAACCCGTCGCCTTGCCTAGGTAATC-3’. The exchange of the fluorophore in delCMV-2xRaf-GBD-mCherry to delCMV-2xRaf-GBD-mCitrine was performed by restriction digest and ligation using AgeI and BsrGI. The perturbation construct mTurquoise2-NES-eDHFR-Rap1a G12V containing the eDHFR dimerization domain and a nuclear export sequence (NES) was generated as follows: First, delCMV-MCP-mCitrine-Rap1a(wt) was generated by restriction digestion and ligation of EYFP-C1 Rap1a(wt) (kind gift from Philippe Bastiaens, MPI Dortmund) and delCMV-MCP-mCitrine (based on MCP-YFP, from Addgene #101160 and delCMV-mCitrine; described in Master thesis Olga Just, TU Dortmund) using BsrGI and MfeI. The dominant positive mutation G12V was then introduced into this construct via site-directed mutagenesis using the primers 5’- GGTCCTTGGTTCAGTAGGCGTTGGGAAG-3’and 5’- CTTCCCAACGCCTACTGAACCAAGGACC-3’ to obtain delCMV-MCP-mCitrine-Rap1aG12V. Finally, mTurquoise2-NES-eDHFR-Rap1a G12V was generated by restriction-free Gibson assembly after PCR amplification of delCMV-MCP-mCitrine-Rap1aG12V using primers 5’-CTCTAGATCCATGCGTGAGTACAAGCTAG-3’ and 5’- AATTGAGTTATTCCACTGGTGTTTTCCTATTTATC-3’, and of mTurquoise2-NES- eDHFR-RhoAQ63L (Kamps et al., 2020), using primers 5’- ACCAGTGGAATAACTCAATTGTTGTTGTTAAC-3’ and 5’-ACTCACGCATGGATCTAGAGGTGGATCC-3’).

### qPCR

Evaluation of siRNA mediated mRNA knockdown was performed using qPCR. PPIB was used as a housekeeping gene to normalize measurements for individual mRNAs and scrambled siRNA was used as a control treatment to calculate knockdown efficiency. qPCR reactions were performed using the “GoTaq^®^ qPCR System” Kit (Promega), using 300 nM of each primer. Primer sequences were selected based on previously published studies: Rap1A (Lin et al., 2015): TGTCTCACTGCACCTTCAATGGCAT (fw), ACGCCTCCTGAACCAAGGACCA (rv); PPIB (Nazet et al., 2019): TTCCATCGTGTAATCAAGGACTTC (fw), GCTCACCGTAGATGCTCTTTC (rv).

### Microscopy

TIRF microscopy was performed using an Olympus IX-81 microscope, equipped with a TIRF-MITICO motorized TIRF illumination combiner, an Apo TIRF 60x/1.45 NA oil immersion objective and a ZDC autofocus device. All TIRF measurements employed a single dichroic mirror (ZT405-440/514/561) that was used in combination with a matched emission filter set (HC 435/40, HC 472/30, HC 542/27 and HC 629/53), a 514 nm OBIS diode laser (150 mW) (Coherent, Inc., Santa Clara, USA), and the Cell R diode lasers (Olympus) with wavelength 405 nm (50 mW), 445 nm (50 mW) and 561 nm (100 mW). In some experiments, TIRF measurements were combined with wide-field illumination via the Spectra X light engine (Lumencor). For detection, an EMCCD camera (C9100-13; Hamamatsu, Herrsching am Ammersee, Germany) was used at medium gain without binning. In addition, the microscope was equipped with a temperature-controlled incubation chamber. Time-lapse live-cell microscopy experiments were carried out at 37 °C in CO2-independent HEPES-stabilized imaging medium (PAN Biotech or Sigma-Aldrich Chemie GmbH/Merck) supplemented with 10% FBS. Automated scanning for cell morphometric analysis was performed using the same microscope using wide-field illumination and an UPlanSApo 10x/0.4 NA air objective and software-based automated focusing.

#### Analysis of cell morphodynamics and local fluorescence signals at the cell edge

Cell morphodynamics were measured and analyzed essentially as described previously (Nanda et al., 2023). Briefly, the local enrichment of the Ras and Rap GTPase activity was investigated in A431 cells by transfecting the corresponding activity sensors together with soluble fluorescent proteins that act as cell volume markers. Movements of the cell edge were analyzed automatically on the basis of the cell volume markers independent of varying GTPase activity signals. The subsequent quantitative analysis procedure was also described previously (Nanda et al., 2023). Briefly, a modified version of the ADAPT plugin (Barry et al., 2015) was combined with custom ImageJ analysis scripts to generate the signal/velocity crosscorrelation functions and protrusion/retraction enrichment measurements. Cell edge movements > 0.5 µm/min were considered to be protrusions, and movements < -0.5 µm/min were considered to be retractions.

#### Analysis of Rac/Rap GTPase activity crosstalk via optogenetic perturbations

Crosstalk between Rac and Rap activity was performed essentially as described before (Nanda et al., 2023). Briefly, photo-activatable Rac1 (mCerulean-PA-Rac1) was co-transfected with the Rap activity sensor delCMV-mCitrine-2xRalGDS-GBD. Light-based activation of Rac1 was performed by TIRF illumination using the 445 nm Cell R diode laser. To prevent excess PA-Rac1 activation, a 10000x neutral-density filter was added into the 445 nm TIRF illumination light path. The built-in neutral-density filter wheel of the microscopy setup was additionally set to 30%. Within photoactivation time intervals, 445 nm TIRF illumination was constantly on, except for the exposure times during image acquisition. Detection of mCerulean was always performed after the experiment. The subsequent analysis was also described previously (Nanda et al., 2023). Briefly, the activity sensor measurement *A_RAP_* was calculated by first measuring the fluorescence intensity of the Rap activity sensor *I_RAP_* via TIRF microscopy in the entire cell adhesion area. These raw intensity measurements were normalized by subtracting the background signal outside the cell area *I_RAP,BG_*, and dividing by the initial, background-corrected intensity value *I_RAP_*_,0_ − *I_RAP_*_,0,*BG*_ before the perturbation:

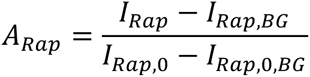

### Analysis of Rap/Rac GTPase activity crosstalk via photochemically-induced dimerization

Crosstalk between Rap and Rac activity was performed using photochemically-induced dimerization, essentially as described previously (Chen et al., 2017). Briefly, local photouncaging of the NvocTMP-Cl dimerizer was performed by a single 200ms illumination pulse (405nm, 180 nW) at a diffraction-limited spot via the FRAP mode of the TIR-MITICO motorized TIRF illumination combiner. Laser power was measured at the aperture of the 60x objective. Stabilized and background-corrected time series of the perturbation and response signals were measured at the perturbation site. To exclude cells in subsequent analyses that did not generate a sufficiently strong perturbation, only cells with ≥ 5% of signal increase in the perturbation signal within 30-50 seconds after illumination were considered.

### Analysis of cell morphology

96 h after siRNA transfection, 15.000 A431 cells per condition were reseeded onto LabTek wells that were previously coated with fibronectin (10 mg/mL). 8 hours after seeding, the cells are washed 3x with 37°C PBS and fixed with prewarmed formaldehyde (3.7% in PBS) for 20 min at 37°C. After 3x washing with PBS, the cells were permeabilized with Triton X-100 (0.1 % in PBS) for 15 min at RT. After 3x washing with PBS, cells were stained for F-actin and nuclei using Rhodamin Phalloidin (1:1000 in PBS) and Hoechst 33342 (1:1000 in PBS) for 30 min at RT in the dark. Cells were washed 3x with PBS and stored at 4°C until imaging within 24 hours of fixing. 64 images were obtained for each condition by automated scanning of individual wells. Subsequently, images were analyzed using Cell Profiler (Carpenter et al., 2006) to measure the cell adhesion area. Briefly, nuclei were detected as primary objects using Hoechst signals and adaptive Otsu thresholding with two classes and a typical object diameter of 5-25 pixels. Then, cells were detected using phalloidin signals as secondary objects using nuclei as input objects, the propagation method and global three class Otsu thresholding. The cell adhesion area was then obtained as the area of the secondary cell objects.

### Software for image and video analysis

Image and video analysis was performed using FIJI (https://imagej.net/software/fiji/), the FIJI Kymograph plugin, the image stabilizer plugin (K. Li, “The image stabilizer plugin for ImageJ,” https://www.cs.cmu.edu/~kangli/code/Image_Stabilizer.html, February, 2008), and a modified version of the ADAPT plugin (Barry et al., 2015; Nanda et al., 2023). Cell Profiler (Carpenter et al., 2006) was used for cell adhesion area calculation, and data were further processed using MatLab. Data plotting and statistical analyses were performed using Prism (GraphPad).

## Acknowledgements

We thank Sven Müller (MPI Dortmund) for expert microscopy support and Philippe Bastiaens (MPI Dortmund) for departmental support and helpful discussions. We also would like to thank Carolin Gierse and Arya Sachan for support in video analyses and helpful discussions. This work was supported by the Deutsche Forschungsgesellschaft DFG project grant 823/9-1 and DFG Principal Investigator grant DE 823/10-1 to L.D.

## Author contributions

L.D., R.L., S.N., H.T.C. designed the research. R.L. performed, analyzed, and optimized the majority of experiments. S.N. developed and characterized the activity sensors and performed initial Rap and Ras activity measurements. H.T.C. performed initial dual Rap/Rac activity measurements. L.D. supervised experiments. L.D. and R.L. wrote the majority of the manuscript. All authors contributed to discussions and manuscript preparation.

## Competing interests

The authors declare no competing interests.

## Description of Additional Supplementary Files

**Movie 1: Spatio-temporal activity paterns of the small GTPases Rap, and Ras via in A431 cells (related to Fig. 1b-c).** Time-lapse TIRF video of the Rap (mCitrine-2xRalGDS-GBD) and Ras (2xRaf-GBD-mCherry) activity sensors. Images were collected with a frame rate of 12/min.

**Movie 2: Spatio-temporal activity paterns of the small GTPases Rap in a migrating A431 cell (related to Fig. 1e).** Time-lapse TIRF video of the Rap activity sensor and a volume marker. Images were collected with a frame rate of 6/min.

**Movie 3: Spatio-temporal activity paterns of the small GTPases Rap, Ras and Rac in A431 cells (related to Fig. 2a,d,g,j).** Time-lapse TIRF videos of the Rap,Ras, Rac or control activity sensors together with corresponding volume markers. Images were collected with a frame rate of 6/min.

**Movie 4: Spatio-temporal activity paterns of the small GTPases Rap and Rac in a migrating A431 cell (related to Fig. 3a).** Time-lapse TIRF video of the Rap (mCitrine-2xRalGDS-GBD) and Rac (mCherry-3xp67phox-GBD) activity sensors in a A431 cell. Images were collected with a frame rate of 30/min.

**Movie 5: Measurement of Rap/Rac crosstalk in A431 cells (related to Fig. 3c).** Top: Time-lapse TIRF videos of the Rap1a perturbation and the Rac activity response in A431 cells. Images were collected with a frame rate of 6/min. Bottom: Intensity plots corresponding to the signals inside the yellow circle in videos above. The y axis corresponds to signals (arbitrary units) and the x axis corresonds to frames in the video. A vertical blue line in the intensity plots indicates the current frame shown above.

**Movie 6: Measurement of Rac/Rap crosstalk in A431 cells (related to Fig. 3f).** Time-lapse TIRF video of the Rac activity response in A431 cells expressing photocativatable Rac (PA-Rac1). Images were collected with a frame rate of 12/min.

## Notes

### Competing Interest Statement

The authors have declared no competing interest.

## References

Arrieumerlou, C., & Meyer, T. (2005). A local coupling model and compass parameter for eukaryotic chemotaxis. Dev Cell, 8(2), 215–227. 10.1016/j.devcel.2004.12.007

Barry, D. J., Durkin, C. H., Abella, J. V, & Way, M. (2015). Open source software for quantification of cell migration, protrusions, and fluorescence intensities. J Cell Biol, 209(1), 163–180. 10.1083/jcb.201501081

Bement, W. M., Leda, M., Moe, A. M., Kita, A. M., Larson, M. E., Golding, A. E., Pfeuti, C., Su, K. C., Miller, A. L., Goryachev, A. B., & von Dassow, G. (2015). Activator-inhibitor coupling between Rho signalling and actin assembly makes the cell cortex an excitable medium. Nat Cell Biol, 17(11), 1471–1483. 10.1038/ncb3251

Boettner, B., & Van Aelst, L. (2009). Control of cell adhesion dynamics by Rap1 signaling. Curr Opin Cell Biol, 21(5), 684–693. 10.1016/j.ceb.2009.06.004

Carpenter, A. E., Jones, T. R., Lamprecht, M. R., Clarke, C., Kang, I. H., Friman, O., Guertin, D. A., Chang, J. H., Lindquist, R. A., Moffat, J., Golland, P., & Sabatini, D. M. (2006). CellProfiler: image analysis software for identifying and quantifying cell phenotypes. Genome Biol, 7(10), R100. 10.1186/gb-2006-7-10-r100

Chen, X., Venkatachalapathy, M., Kamps, D., Weigel, S., Kumar, R., Orlich, M., Garrecht, R., Hirtz, M., Niemeyer, C. M., Wu, Y. W., & Dehmelt, L. (2017). “Molecular Activity Painting”: Switch-like, Light-Controlled Perturbations inside Living Cells. Angew Chem Int Ed Engl, 56(21), 5916–5920. 10.1002/anie.201611432

Devreotes, P. N., Bhattacharya, S., Edwards, M., Iglesias, P. A., Lampert, T., & Miao, Y. (2017). Excitable Signal Transduction Networks in Directed Cell Migration. Annu Rev Cell Dev Biol, 33, 103–125. 10.1146/annurev-cellbio-100616-060739

Diekmann, D., Abo, A., Johnston, C., Segal, A. W., & Hall, A. (1994). Interaction of Rac with p67phox and regulation of phagocytic NADPH oxidase activity. *Science (New York*, N.Y.), 265(5171), 531–533. 10.1126/SCIENCE.8036496

Ferro, E., Goitre, L., Baldini, E., Retta, S. F., & Trabalzini, L. (2014). Ras GTPases are both regulators and effectors of redox agents. Methods in Molecular Biology (Clifton, N.J.), 1120, 55–74. 10.1007/978-1-62703-791-4_5

Graessl, M., Koch, J., Calderon, A., Kamps, D., Banerjee, S., Mazel, T., Schulze, N., Jungkurth, J. K., Patwardhan, R., Solouk, D., Hampe, N., Hoffmann, B., Dehmelt, L., & Nalbant, P. (2017). An excitable Rho GTPase signaling network generates dynamic subcellular contraction patterns. J Cell Biol, 216(12), 4271– 4285. 10.1083/jcb.201706052

Guilluy, C., Garcia-Mata, R., & Burridge, K. (2011). Rho protein crosstalk: another social network? Trends Cell Biol, 21(12), 718–726. 10.1016/j.tcb.2011.08.002

Hobbs, G. A., Der, C. J., & Rossman, K. L. (2016). RAS isoforms and mutations in cancer at a glance. Journal of Cell Science, 129(7), 1287–1292. 10.1242/JCS.182873/260209/AM/RAS-ISOFORMS-AND-MUTATIONS-IN-CANCER-AT-A-GLANCE

Joshi, M. S., Stanoev, A., Huebinger, J., Soetje, B., Zorina, V., Roßmannek, L., Michel, K., Müller, S. A., & Bastiaens, P. I. (2023). The EGFR phosphatase RPTPγ is a redox-regulated suppressor of promigratory signaling. The EMBO Journal, 42(10). 10.15252/EMBJ.2022111806

Kamps, D., Koch, J., Juma, V. O., Campillo-Funollet, E., Graessl, M., Banerjee, S., Mazel, T., Chen, X., Wu, Y. W., Portet, S., Madzvamuse, A., Nalbant, P., & Dehmelt, L. (2020). Optogenetic Tuning Reveals Rho Amplification-Dependent Dynamics of a Cell Contraction Signal Network. Cell Rep, 33(9), 108467. 10.1016/j.celrep.2020.108467

Klijn, C., Durinck, S., Stawiski, E. W., Haverty, P. M., Jiang, Z., Liu, H., Degenhardt, J., Mayba, O., Gnad, F., Liu, J., Pau, G., Reeder, J., Cao, Y., Mukhyala, K., Selvaraj, S. K., Yu, M., Zynda, G. J., Brauer, M. J., Wu, T. D., … Zhang, Z. (2014). A comprehensive transcriptional portrait of human cancer cell lines. Nature Biotechnology 2014 33:3, 33(3), 306–312. 10.1038/nbt.3080

Lafuente, E. M., van Puijenbroek, A. A. F. L., Krause, M., Carman, C. V., Freeman, G. J., Berezovskaya, A., Constantine, E., Springer, T. A., Gertler, F. B., & Boussiotis, V. A. (2004). RIAM, an Ena/VASP and Profilin ligand, interacts with Rap1-GTP and mediates Rap1-induced adhesion. Developmental Cell, 7(4), 585–595. 10.1016/J.DEVCEL.2004.07.021

Lambert, J. M., Lambert, Q. T., Reuther, G. W., Malliri, A., Siderovski, D. P., Sondek, J., Collard, J. G., & Der, C. J. (2002). Tiam1 mediates Ras activation of Rac by a PI(3)K-independent mechanism. Nature Cell Biology, 4(8), 621–625. 10.1038/NCB833

Lin, K. T., Yeh, Y. M., Chuang, C. M., Yang, S. Y., Chang, J. W., Sun, S. P., Wang, Y. S., Chao, K. C., & Wang, L. H. (2015). Glucocorticoids mediate induction of microRNA-708 to suppress ovarian cancer metastasis through targeting Rap1B. Nature Communications 2015 6:1, 6(1), 1–13. 10.1038/ncomms6917

Miao, Y., Bhattacharya, S., Edwards, M., Cai, H., Inoue, T., Iglesias, P. A., & Devreotes, P. N. (2017). Altering the threshold of an excitable signal transduction network changes cell migratory modes. Nat Cell Biol, 19(4), 329–340. 10.1038/ncb3495

Nanda, S., Calderon, A., Sachan, A., Duong, T. T., Koch, J., Xin, X., Solouk-Stahlberg, D., Wu, Y. W., Nalbant, P., & Dehmelt, L. (2023). Rho GTPase activity crosstalk mediated by Arhgef11 and Arhgef12 coordinates cell protrusion-retraction cycles. Nature Communications 2023 14:1, 14(1), 1–17. 10.1038/s41467-023-43875-y

Nazet, U., Schröder, A., Grässel, S., Muschter, D., Proff, P., & Kirschneck, C. (2019). Housekeeping gene validation for RT-qPCR studies on synovial fibroblasts derived from healthy and osteoarthritic patients with focus on mechanical loading. PLOS ONE, 14(12), e0225790. 10.1371/JOURNAL.PONE.0225790

Patwardhan, R., Nanda, S., Wagner, J., Stockter, T., Dehmelt, L., & Nalbant, P. (2024). Cdc42 activity in the trailing edge is required for persistent directional migration of keratinocytes. Molecular Biology of the Cell, 35(1). 10.1091/MBC.E23-08-0318/MC-E23-08-0318-S09.MOV

Ridley, A. J. (2001). Rho GTPases and cell migration. Journal of Cell Science, 114(Pt 15), 2713–2722. 10.1242/JCS.114.15.2713

Ridley, A. J., Paterson, H. F., Johnston, C. L., Diekmann, D., & Hall, A. (1992). The small GTP-binding protein rac regulates growth factor-induced membrane ruffling. Cell, 70(3), 401–410. https://www.ncbi.nlm.nih.gov/pubmed/1643658

Scarpa, E., & Mayor, R. (2016). Collective cell migration in development. The Journal of Cell Biology, 212(2), 143–155. 10.1083/JCB.201508047

Steffen, A., Ladwein, M., Dimchev, G. A., Hein, A., Schwenkmezger, L., Arens, S., Ladwein, K. I., Margit Holleboom, J., Schur, F., Victor Small, J., Schwarz, J., Gerhard, R., Faix, J., Stradal, T. E., Brakebusch, C., & Rottner, K. (2013). Rac function is crucial for cell migration but is not required for spreading and focal adhesion formation. J Cell Sci, 126(Pt 20), 4572–4588. 10.1242/jcs.118232

Svitkina, T. M., Verkhovsky, A. B., McQuade, K. M., & Borisy, G. G. (1997). Analysis of the actin-myosin II system in fish epidermal keratocytes: mechanism of cell body translocation. J Cell Biol, 139(2), 397–415. https://www.ncbi.nlm.nih.gov/pubmed/9334344

Trepat, X., Chen, Z., & Jacobson, K. (2012). Cell migration. Comprehensive Physiology, 2(4), 2369–2392. 10.1002/CPHY.C110012

Watanabe, N., & Mitchison, T. J. (2002). Single-molecule speckle analysis of actin filament turnover in lamellipodia. Science, 295(5557), 1083–1086. 10.1126/science.1067470 295/5557/1083 [pii]

Wittchen, E. S., Van Buul, J. D., Burridge, K., & Worthylake, R. A. (2005). Trading spaces: Rap, Rac, and Rho as architects of transendothelial migration. Current Opinion in Hematology, 12(1), 14–21. 10.1097/01.MOH.0000147892.83713.A7

Wu, Y. I., Frey, D., Lungu, O. I., Jaehrig, A., Schlichting, I., Kuhlman, B., & Hahn, K. M. (2009). A genetically encoded photoactivatable Rac controls the motility of living cells. Nature, 461(7260), 104–108. nature08241 [pii] 10.1038/nature08241

Yang, H. W., Collins, S. R., & Meyer, T. (2016). Locally excitable Cdc42 signals steer cells during chemotaxis. Nat Cell Biol, 18(2), 191–201. 10.1038/ncb3292

